# Isoflurane reduces feedback in the fruit fly brain

**DOI:** 10.1101/161976

**Authors:** Dror Cohen, Bruno van Swinderen, Naotsugu Tsuchiya

**Affiliations:** School of Psychological Sciences, Monash University, Melbourne, Australia, 3168; Queensland Brain Institute,The University of Queensland, Brisbane, Australia, 4072; Monash Institute of Cognitive and Clinical Neuroscience, Monash University, Australia, 3168

## Abstract

Hierarchically organized brains communicate through feedforward and feedback pathways. In mammals, feedforward and feedback are mediated by higher and lower frequencies during wakefulness. Feedback is preferentially impaired by general anesthetics. This suggests feedback serves critical functions in waking brains. The brain of *Drosophila melanogaster* (fruit fly) is also hierarchically organized, but the presence of feedback in these brains is not established. Here we studied feedback in the fruit fly brain, by simultaneously recording local field potentials (LFPs) from low-order peripheral structures and higher-order central structures. Directed connectivity analysis revealed that low frequencies (0.1-5Hz) mediated feedback from the center to the periphery, while higher frequencies (10-45Hz) mediated feedforward in the opposite direction. Further, isoflurane anesthesia preferentially reduced feedback. Our results imply that similar spectral characteristics of feedforward and feedback may be a signature of hierarchically organized brains and that general anesthetics may induce unresponsiveness by targeting the mechanisms that support feedback.

## Manuscript

Many complex networks including brains are hierarchically organized. In hierarchical systems information travels both from the bottom to the top of the hierarchy, known as feedforward (FF), and from the top to the bottom of the hierarchy, known as feedback (FB). It is only recently that the dynamic characteristics of neural FB and FF have been reported. For example, studies that investigated directed connectivity using Granger causality across the awake monkey visual hierarchy have shown that the alpha/beta and gamma bands preferentially mediate FB and FF respectively (*1, 2*). This pattern of low frequency FB/high frequency FF has also been reported across the visual (*3*) and auditory system (*4*) in awake humans. These studies imply that FB is mediated by lower frequencies than those that mediate FF. One potential explanation for these observations is that the slower time scale of FB reflect the incorporation of memory and expectation for the modulation of quickly changing sensory inputs (FF) (*5*).

A separate line of inquiry suggests that FB in particular is crucial for waking brain functions. A number of studies have demonstrated that general anesthetics preferentially reduce FB from frontal to posterior areas as measured using human scalp level electroencephalography EEG (*6-11*) (but see (*12-14*)

Here we investigated the dynamic characteristics of feedback and feedforward in the small brains of fruit flies. Anatomically, fly brains are hierarchically organized from peripheral sensory layers to central nuclei, which are responsible for more abstract computation (*15-17*). Like mammals, wakeful insect brains also face functional challenges in adjusting fast sensory inputs based on prior experience on a slower time scale (*18-21*). Thus, it is plausible that similar dynamic characteristics of FF and FB processing will be present in fruit flies.

We analyzed local field potentials (LFPs) recorded from the brains of behaving fruit flies (see Figure 1 and Figure 1S for more details). LFPs reflect aggregate neural activity and have been shown to relate to various sensory and cognitive processing in flies (*22-27*). Tethered flies were placed on an air supported ball and a linear electrode array was inserted laterally across one hemisphere of their brains (Figure 1a, b). This recording technique combined with bipolar rereferencing (see Local field potential pre-processing in supporting material) provides a set of 14 LFP signals from both the periphery and higher order structures situated more centrally in the fly brain (Figure 1c).

**Figure 1.**
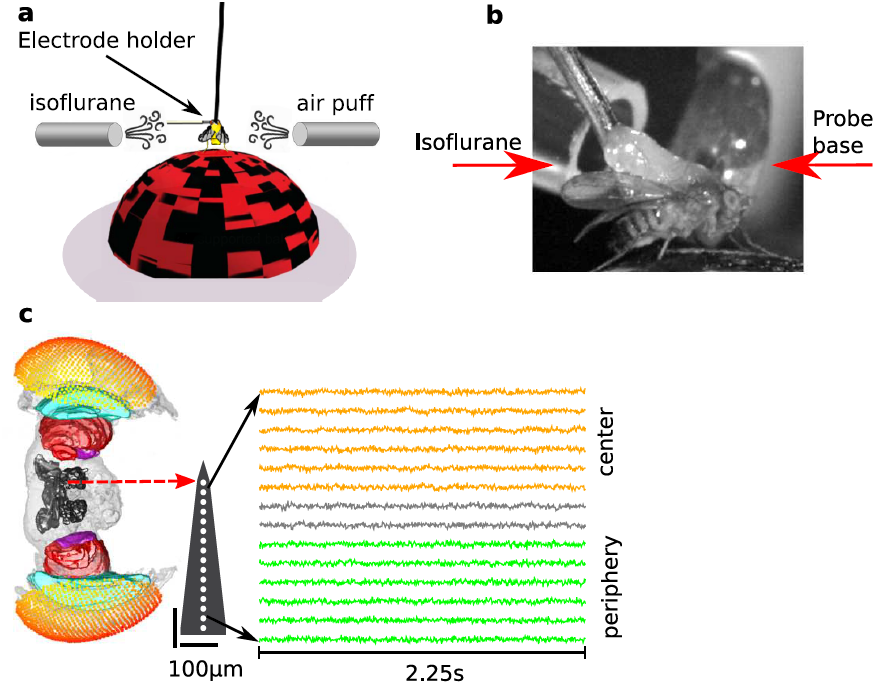
Recording LFPs from the center and periphery of the fruit fly brain. **a)** Flies were dorsally fixed to a tungsten rod and placed on an air-supported ball where they could freely walk. Isoflurane in different volumetric concentrations was delivered through a rubber hose. Air puffs were used to gauge flies’ responsiveness. 16-contact electrode probe mounted on an electrode holder was inserted laterally from the left into the fly’s eye. Only the electrode holder is visible at the depicted scale. **b)** A close-up view contralateral to the electrode insertion site showing the tethered fly, isoflurane delivery hose, and probe base. **c)** Example of pre-processed (see supporting material) LFPs data before anesthesia (0% isoflurane) from one fly. A standardized fly brain is shown for comparison (*22, 23*). The electrode contacts are indicated by white dots (not to scale). LFPs were subsampled to 250Hz to aid visualization. Six (pre-processed) channels are grouped as peripheral, estimated to correspond to the optic lobe (green), and another six channels are grouped as central, estimated to correspond to the central brain (orange).

Low frequency power in the center was greater than in the periphery, demonstrating that neural activity is distinguishable across these two anatomical areas (Figure 2a). Coherence between the center and periphery was highest but most variable for low frequencies and decreased in value as frequency increased. Because coherence is confined to 0-1 it is inherently biased and care must be taken when assessing low coherence values. By empirically estimating the coherence bias in our data (see supplemental material) we found that coherence values were above bias for 0.1-45Hz (Figure 2b). This assures that for 0.1-45Hz we can detect central-peripheral connectivity at the level of LFPs. To dissect the directionality of this connectivity we used Granger causality (GC) analysis (*28-30*). When we examined the frequency spectra of GC we found that for low frequencies (0.1-5Hz) GC influences from the center to the periphery (FB) appeared greater than GC influences from the periphery to the center (FF), while the reverse appeared true for higher frequencies (10-45Hz) (Figure 2c). As is apparent in Figure 2c, spectral GC influences scaled similarly to coherence (i.e., the lower frequency, the larger the value and variance). To correct for these statistical characteristics we employed a normalized measure known as the directed asymmetry index (DAI) (*1, 3, 9*). The DAI is defined as

**Figure 2.**
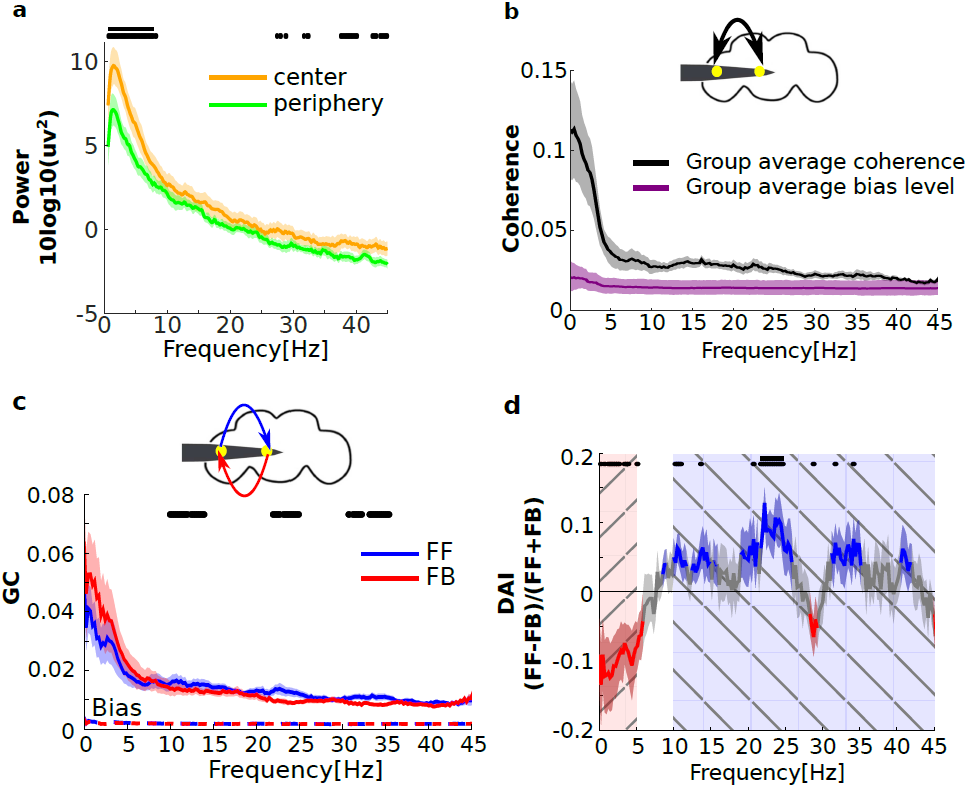
Frequency-specific Granger causal influences in the fruit fly brain. **a)** Group average center (orange) and periphery (green) power in 0% isoflurane (before any anesthesia). Black dots and solid line mark uncorrected and corrected significant differences between central and peripheral channels at the p< 0.05 level respectively (see supporting material). Shaded area represents sem across flies (N=13). **b)** Group average center-periphery coherence in 0% isoflurane (black) is above the bias level (purple) for 0.1-45Hz. Shaded area represents 95% confidence intervals (see supporting material). A schematic representation of the fly head and inserted electrode showing the estimated positions of one central (C) and one peripheral (P) channel is depicted above. **c)** Group level feedforward (FF, blue) and feedback (FB, red) GC influences before any anesthesia (0% isoflurane). Black dots mark significant differences between FF and FB at the p< 0.05 level (uncorrected). The FF and FB GC bias levels (see supporting material) are shown as dashed red and blue lines, which almost completely overlap near 0. Shaded area represents sem across flies N=13). **d)** Group level directed asymmetry index (DAI) before any anesthesia (0% isoflurane). Positive values indicate FF> FB (frequencies for which the DAI is one sem or more below zero are shown in red) and negative values indicate FF< FB (frequencies for which the DAI is one sem or more above zero are shown in blue). Black dots and solid line mark significant uncorrected and corrected differences from zero at the p< 0.05 level. Low (0.1-5Hz) and high (10-45Hz) frequencies are indicated by the red and blue hatchings. Shaded area represents sem across flies (N=13).

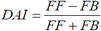

which is positive when FF is stronger than FB and vice versa. For the DAI the variance was uniform across all frequencies (Figure 2d). The DAI was negative for lower frequencies (~ 0.1-5Hz), indicating that FB is stronger than FF. For higher frequencies (10-45Hz) the DAI is predominantly positive, indicating that FF is stronger than FB.

To summarize these results, we averaged the DAI for low (0.1-5Hz) and high (10-45Hz) frequencies separately. We found that in awake flies FB is stronger than FF for low frequencies (0.1-5Hz) (Figure 3a, p< 0.05, group-level permutation tests, see supporting material) and that FB is weaker than FF for high frequencies (10-45Hz) (Figure 3b, p< 0.01). This global pattern of results, that is, the dominance of feedback in lower frequencies and the dominance of feedforward in higher frequencies, is generally consistent with what has been reported in the awake mammalian brain (*1-4*).

**Figure 3.**
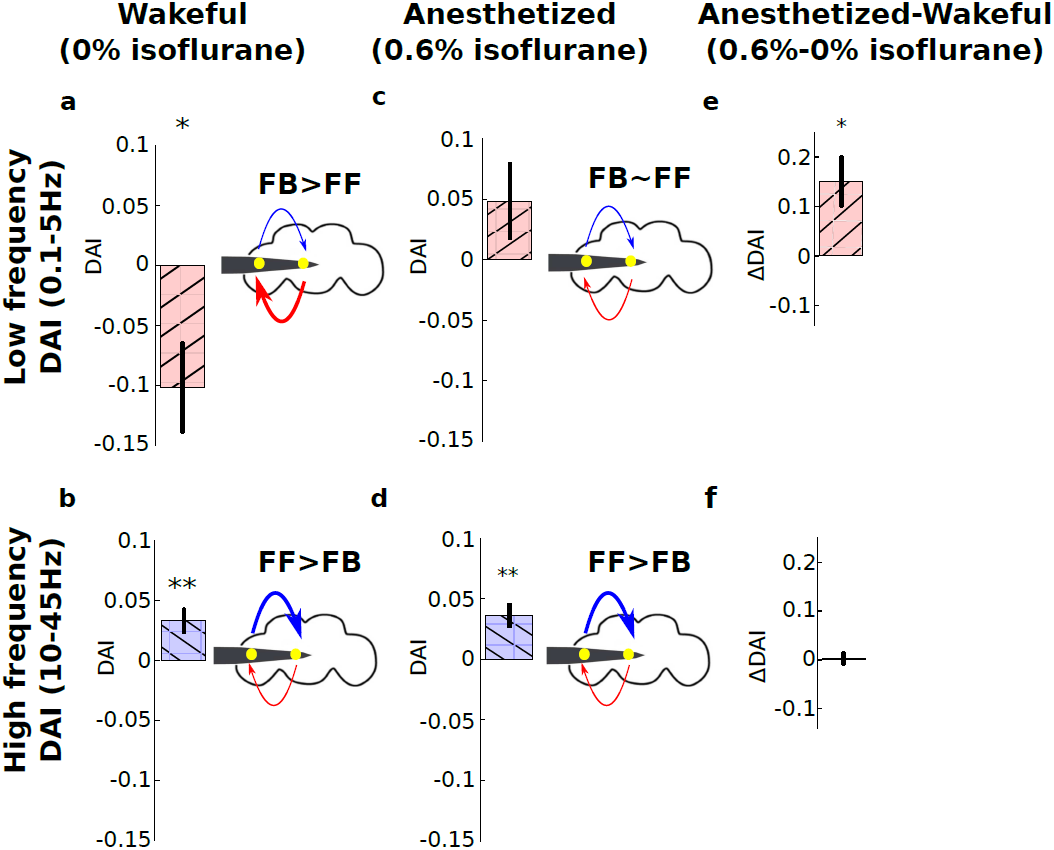
Isoflurane reduces low frequency (0.1A5Hz) feedback in the fruit fly brain. Group level DAI averaged for low (0.1-5Hz) (**a**) and high (10-45Hz) frequencies (**b**) before any anesthesia (0% isoflurane). Negative and positive DAI values indicate that FB> FF for low frequencies and that FB< FF for high frequencies. The fly head schemas above the graphs depict this as thicker FF (red, **a**) and FB (blue, **b**) arrows. **c**-**d)** same as **a** and **b** during 0.6% isoflurane. For low frequencies FB and FF are roughly balanced (**c**), while FF remains stronger than FB for higher frequencies (**d**). The effect of 0.6% isoflurane on low (**e**) and high frequencies (**f**), obtained by subtracting values in 0% isoflurane (**a,b**) from 0.6% isoflurane (**c,d**). ^*^ and ^**^ indicate significant differences from zero at the p< 0.05 and p< 0.01 level respectively (group-level permutation tests, see supporting material). Error bars represent sem across flies N=13.

Next, we examined the relevance of FB and FF to wakefulness by exposing flies to the general anesthetic isoflurane (Figure S1 and supporting material). At a concentration of 0.6% isoflurane, the flies are rendered unresponsive, as previously probed by air puffs (*31*). Isoflurane (0.6%) reduced LFP power in both the center and periphery, indicating an overall reduction in activity over waking levels (Figure S2a). This reduction in spontaneous power was accompanied by a reduction in 10-30Hz coherence, indicating impaired center-periphery connectivity (Figure S2b).

Isoflurane had frequency- and direction-specific effects on GC (Figure 3). In terms of magnitude, isoflurane clearly reduced FF and FB GC values for high frequencies (10-40Hz), whereas reductions in lower frequencies (0.1-5Hz) were more variable (Figure S2c). The DAI, which normalizes the magnitude and variance across frequencies, was significantly greater during anesthesia for low frequencies (Figure S2d). This meant that during anesthesia the dominance of FB over FF was lost (Figure 3c). In contrast, the FF dominance over FB in higher frequencies remained (Figure 3d). When comparing the wakeful and anesthetized conditions (Δ DAI = DAI at isoflurane 0% - DAI at isoflurane 0.6%, Figure 3e and f), the above results were confirmed as significantly positive Δ DAI for low (0.1-5Hz, p< 0.05) but not for high frequencies (10-45Hz, p=0.8).

What can explain our finding that FB and FF are mediated by lower (0.1-5Hz) and higher (10-40Hz) frequencies, respectively? From a predictive coding view, low frequency FB may reflect the slower time scales required to form ‘predictions’ while higher frequency FF reflects the faster time scales in which ‘prediction errors’ are processed (*5, 32, 33*). Thus, our results are consistent with a predictive coding interpretation of brain function in fruit flies. Indeed, a number of recent studies have demonstrated predictive behaviors in insects and begun to investigate their neural correlates (*18-21*). Our finding that isoflurane preferentially reduced low frequency FB suggests that general anesthetics impair top down predictions rather than bottom up prediction errors, and that arousal and waking awareness depend primarily on the former.

## Acknowledgments

We would like to thank Jakob Hohwy for useful comments on this manuscript.

## Materials and Methods

### Overview

We analyzed local field potentials (LFPs) data recorded from the brains of awake and anesthetized fruit flies (*1*). In this paper we report a part of the data not reported in (*1*). Here we focus on Granger causality of spontaneous LFPs, and how this is affected by isoflurane. In this section, we briefly recap the experimental setup. For the full experimental details see (*1*).

Thirteen female laboratoryEreared *D. melanogaster* (Canton S wild type) flies (3–7 d post eclosion) were collected under cold anesthesia, tethered and positioned on an airEsupported Styrofoam ball. Linear silicon probes with 16 electrodes separated by 25um (Neuronexus 3mmE25E177) were inserted laterally to the eye of the fly until the most peripheral electrode site was just outside the eye. This probe covers approximately half of the fruit fly brain and records neural activity from both peripheral and central brain structures (Figure 1c). A fine tungsten wire was inserted in the thorax and used as a reference electrode.

Isoflurane was delivered through a rubber hose connected to an evaporator (Figure 1a,b). We tested the effects of isoflurane at concentrations of 0% and 0.6% (for details see (*1*)). An olfactory stimulus controller was used to deliver six air puffs to gauge the behavioral responsiveness of the flies in each concentration of isoflurane. Fly movement activity in response to the air puff was recorded with a camera. The video data was used to confirm that 0.6% isoflurane abolished the behavioural responsiveness and that behavioural responsiveness returned after recovery (*1*).

An experiment consisted of several blocks, each at a different concentration of isoflurane. Each block started with the delivery of a series of air puffs (used to gauge the behavioural responsiveness), followed by approximately 18s of rest, which is the critical period of data for this paper. Visual stimuli were presented, lasting 248s in total, followed by an additional 30s of rest. A series of air puffs were delivered again and then the isoflurane concentration was changed. After 180s of adjustment to the new isoflurane concentration the experimental block was repeated (Figure S1).

In (*1*) we focused on the analysis of the visuallyEevoked data (i.e., data corresponding to the 248s period of visual stimuli presentation). Here we focused on an 18s period before the presentation of visual stimuli (Figure S1).

This period is better suited for coherence and Granger causality than the 248s period in which we presented visual stimuli because the presentation of stimuli can result in erroneous interpretation of coherence and Granger causality (*2, 3*). We focused on the period before rather than after visual stimuli presentation because we reasoned the latter period may also reflect residual visual activity.

### Local field potential pre-processing

LFPs were recorded at 25kHz, downsampled to 1000 Hz and the most peripheral electrode site was removed from the analysis. The remaining 15 unipolar channels were bipolar rereferenced by subtracting adjacent channels to obtain a set of 14 differential signals which we refer to simply as ‘channels’ (*4, 5*).

The 18s period was divided into 8 consecutive epochs of 2.25s. Line noise was removed from each epoch and bipolar rereferenced channel using three tapers, a window of 0.75s and a step size of 0.375s using the rmlinesmovingwinc.m function from the Chronux toolbox (http://chronux.org/;(*6*)). Each epoch and bipolar rereferenced channel was linearly detrended and zEscored by removing the temporal mean and dividing by the temporal standard deviation (*4, 7*). Figure 1c shows example of the resulting LFPs from one fly.

### Analyzing power

For power analysis we linearly detrended the preEprocessed LFPs without the zEscoring operation, because zEscoring obscures overall shifts in power when comparing preE and postEanesthesia (Figure S2a). We calculated power for each preEprocessed channel *i* (i =[1-14]), epoch *e* (1-8), *k%* isoflurane concentration (k=[0 (air), 0.6]) and frequency ω, 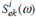 over the 2.25s epochs. Using 3 tapers, the half bandwidth was 0.89Hz (*8*). We further averaged power across 8 epochs to obtain one estimate of power for each channel and isoflurane concentration 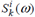.

We analyzed the average power across channels within the center 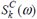 and periphery 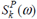 in units of 10log10μ V^2^, following the division of channels into peripheral channels=[1–6] and central channels=[9–14] in (*1*)

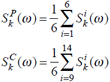

Center and periphery power in 0% isoflurane is shown in Figure 2a.

We report the effect of isoflurane on center power as the difference in power between 0.6% and 0% in decibels (dB)

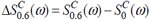

Analogous derivations give the effect of 0.6% isoflurane on periphery power 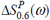 The effect of isoflurane on center and periphery power is shown in Figure S2a.

Previous work has shown *that Drosophila* heartbeat can manifest as low frequency (2-4Hz) oscillatory activity in central channels (*9*). Heartbeats are unlikely to affect the results presented here because we use bipolar rereferencing which eliminates common input to neighboring channels, as may be expected from a muscle artifact. To further rule out heartbeat as a potential confound we reanalyzed Granger causality in our data after excluding three flies in which we detected potential heartbeat in central channels by visual inspection of the power spectrums. The results of this analysis closely matched those obtained with all thirteen flies, indicating the any heartbeat contamination is unlikely to affect our results.

### Coherence analysis

Coherence measures the strength of linear dependency between two signals. It is defined as

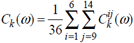

where *CS*^*ij*^ (*ω*)is the crossEspectrum between channels i and j, and *S*^*i*^ (*ω*) and *S* ^*j*^ (*ω*) are the power spectra of channels i and j respectively. To estimate coherence for each bipolar rereferenced channel pair *i* and *j* (i,j =[1-14]), epoch *e* (1E8) and *k%* isoflurane concentration (k=[0 (air), 0.6]) we used the multitaper method with 9 tapers for each 2.25s epoch, giving a half bandwidth of 2.22Hz (*8*). The auto and cross spectra for channels *i* and *j* at isoflurane concentration *k*% were averaged across the 8 epochs to give one estimate of coherence at each isoflurane concentration.

We report centerEperiphery coherence as averaged across all centerEperiphery pairs

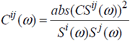

Coherence in 0% isoflurane is shown in Figure 2b. The effect of isoflurane is defined analogously to that of power and reported in Figure S2b.

### Estimating the coherence bias

Because coherence is limited to the 0E1 range it has an inherent positive bias (*10*). We used surrogate data statistics to estimate the empirical coherence bias in our data. The advantages of this method are 1) that it provides an empirical lower bound above which coherence may be considered significant and 2) that it accounts for potentially frequencyEspecific coherence biases sometime observed for two timeEseries with similar power spectrums (*11*).

For each of 13 flies we created surrogate data in which the phase of the signal was randomized but amplitude kept the same (*11, 12*). We performed this 200 times for each channel (1-14), epoch (1-8) and isoflurane concentration ([0%, 0.6%]) resulting in surrogate data with the same power spectra as the original data but with nearEzero cross spectra between channels. The surrogate data describes the null hypothesis that there is no coherence in the data.

To estimate the coherence bias at *k*% isoflurane *C*_*bias,k*_ (*ω*) we calculated coherence as described above for each of the 200 surrogate data sets, and averaged the estimates across all 200 surrogates. We obtained the 95% confidence intervals by determining the 2.5 and 97.5 percentiles across the 200 surrogates for each fly. The group mean coherence bias and the mean 95% confidence intervals across all 13 flies during 0% isoflurane are shown in Figure 2a.

When we compared the coherence bias in 0.6% and 0% isoflurane, we observed differences in low frequencies (0.1-5Hz), but the magnitude of the difference was very small (largest difference was 0.0025 at around 2Hz). Accordingly, our results on coherence in 0.6% and 0% isoflurane with or without bias correction were nearly identical.

### Non-parametric Granger causality analysis

In simple terms, a signal X is said to *Granger-cause* a signal - if past values of X improve predictions of future values of Y. This notion of causality is based on modeling X and Y as autoregressive processes. Granger causality can be estimated in either the time or frequency domain. The latter provides a “spectrum” of GC influences. Examining GC influences at different frequencies is important because recent work shows that different causal interactions are mediated by different frequencies (*4, 13, 14*).

Granger causality can be estimated parametrically by fitting an autoregressive model to the data (*15-17*). The drawback of this approach is that it requires selection of the model order (*18*). Alternatively, grangerEcausality can be nonEparametrically estimated directly from the spectral density matrix (*19, 20*). The spectral density matrix for channels i and j at isoflurane concentration 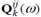 is obtained by setting the diagonal elements to the autoEspectra and the crossEdiagonal elements to the crossEspectra (*20*)

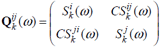

We estimated the spectral density matrix using the same estimates of autoE and crossEspectra as those used for coherence analysis.

Granger-Ecausal influences from channel i to channel j at isoflurane concentration 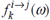 were estimated by factorizing the spectral density matrix using the *sfactorization_wilson*.m and *ft_connectivity_Granger.m* functions from the FieldTrip toolbox (*21*).

### Estimating the Granger causal influences bias

GC influences are always positive and therefore have a positive bias. We checked whether the GC bias differed between FF and FB or across isoflurane concentrations. To estimate the GC bias we used the same procedure for the estimation of the coherence bias. Specifically, we reEestimated FF and FB GC influences using the phase randomized surrogate data as we did for the coherence bias estimation. This procedure results in 200 estimates of FF and FB GC influences for each fly, at each isoflurane concentration *k*%. We obtained the final estimate of the FF and FB GC bias by averaging across all 200 surrogates.

The differences between FF and FB GC biases at each isoflurane concentration, as well as the differences in FF and FB between isoflurane concentrations were negligible. The maximum difference in group average bias was 0.0008 (difference between FF and FB at 0.6% isoflurane at around 5Hz). The difference in GC bias averaged across 0.1E45Hz were all negligible: 0.00006 (between FF and FB in 0% isoflurane), 0.0002 (between FF and FB in 0.6% isoflurane), 0.0002 (between 0% and 0.6% isoflurane for FF) and 0.0001 (between 0% and 0.6% isoflurane for FB). We show the FF and FB GC biases in 0% isoflurane as dotted lines in Figure 2c.

### Grouping feedforward and feedback influences

In the fly brain, visual information enters through the retina and is sequentially processed by the lamina, medulla and lobula, collectively referred to as the optic lobe (*22, 23*). Visual projection neurons (VPNs) convey information from lower to higher-Eorder visual processing centers, called optic glomeruli, which are located more centrally (*24*). Relatively little is known about computations in the optic glomeruli, but recent studies suggest a role in second-Eorder motion processing (*25*) and processing that mediates natural avoidance behaviors (*26*). Specific optic glomeruli, such as the ventrolateral protocereberum (*27*), send projections back to the lobula and medulla, as well as reciprocated projections to other central structures (*24, 28*). Our simple summary of this connectome is that visual information enters in the periphery and sequentially conveyed through FF connections towards more central optic glomeruli, which convey FB back to the optic lobe.

To obtain an overall estimation FF GC influences at k% isoflurane 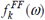 we averaged across all GC influences 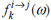from the periphery to the center

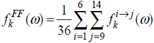

where channel 1 is the most peripheral and channel 14 is the most central. Analogous derivations give FB GC influences

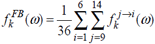

FF and FB GC influences at 0% are shown in Figure 2c.

We report the effect of 0.6% isoflurane on FF GC influences by subtracting values in 0% isoflurane from values in 0.6% isoflurane

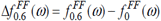

Analogous derivation gives the effect of 0.6% isoflurane on FB GC influences 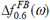.

### Directed Asymmetry Index analysis

Because the magnitude and variance of GC influences are dependent on the frequencies, some form of normalization is necessary for statistical assessment. For this purpose, we adopt the Directed Asymmetry Index (*4, 13, 29*), which is defined as

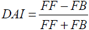

The numerator of the DAI captures the predominant direction of the GC influence. The denominator provides normalization for the total GC influences. DAI allows comparisons across conditions where there may be an overall net increase or decrease in GC influences.

Given our enumeration of channels (i.e. iE> j indicates FF influences if i< j), the DAI between channel i and j at isoflurane concentration k% is given by

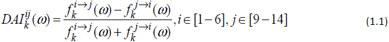

We report the DAI averaged over all centerEperiphery channel pairs

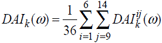

### Statistical analysis

To test for statistical differences between 0% and 0.6% isoflurane across frequencies (power, coherence, GC and DAI spectrums) we used randomizationEbased, nonEparametric statistical tests with clusterEbased multiple comparison correction (*30*). Each of the 13 flies provided one spectrum for each condition (0% and 0.6% isoflurane). We then performed a paired t tests across 13 flies at each frequency to obtain t scores. If we found a cluster of continuous frequencies at p< 0.05 level, then we treated them as a firstElevel significant cluster. As a secondElevel statistics, we used the sum of the t scores across frequencies within each cluster. We then preformed 8191 randomization corresponding to all possible randomization of thirteen flies across the two conditions (2^ 13-1=8191) and calculated the secondElevel statistics for each randomization. From each randomization, the largest secondElevel statistics was chosen and used for constructing a randomized distribution. We compared the observed secondElevel statistics against the randomized distribution to obtain p values. Figure 2a and d and Figure S2a, b, c and d show the results from the firstElevel (uncorrected) t tests as well as the secondElevel clusterEbased statistics, which corrects for multiple comparisons across frequencies. When we compared DAI spectrums against zero (Figure 2d), we randomized the FF and FB labels for each fly (*13*).

For testing whether the DAIs averaged across low and high frequencies were different from zero (Figure 3a&d) we randomized the FF and FB labels and used the group mean as the test metric. For testing the effect of isoflurane on the DAIs averaged across low and high frequencies (Figure 3e&f) we randomized the isoflurane concentrations labels and used the group mean difference between the conditions as the test metric.

---

**Figure S1.**
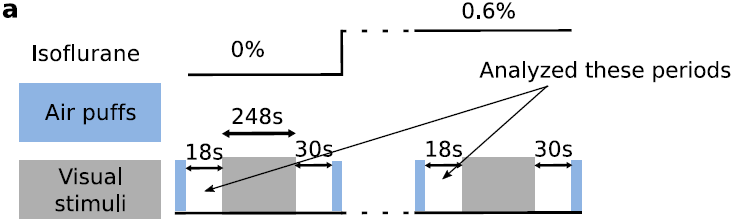
Experimental protocol. An experiment consisted of multiple blocks at different concentration of isoflurane (top black line). Each block proceeded with (1) a series of air puffs (light blue rectangles) followed by 18s of rest; (2) presentation of visual stimuli (248s) (3) 30s of rest followed by a series of air puffs; (4) isoflurane concentration change followed by 180s of rest.

**Figure S2.**
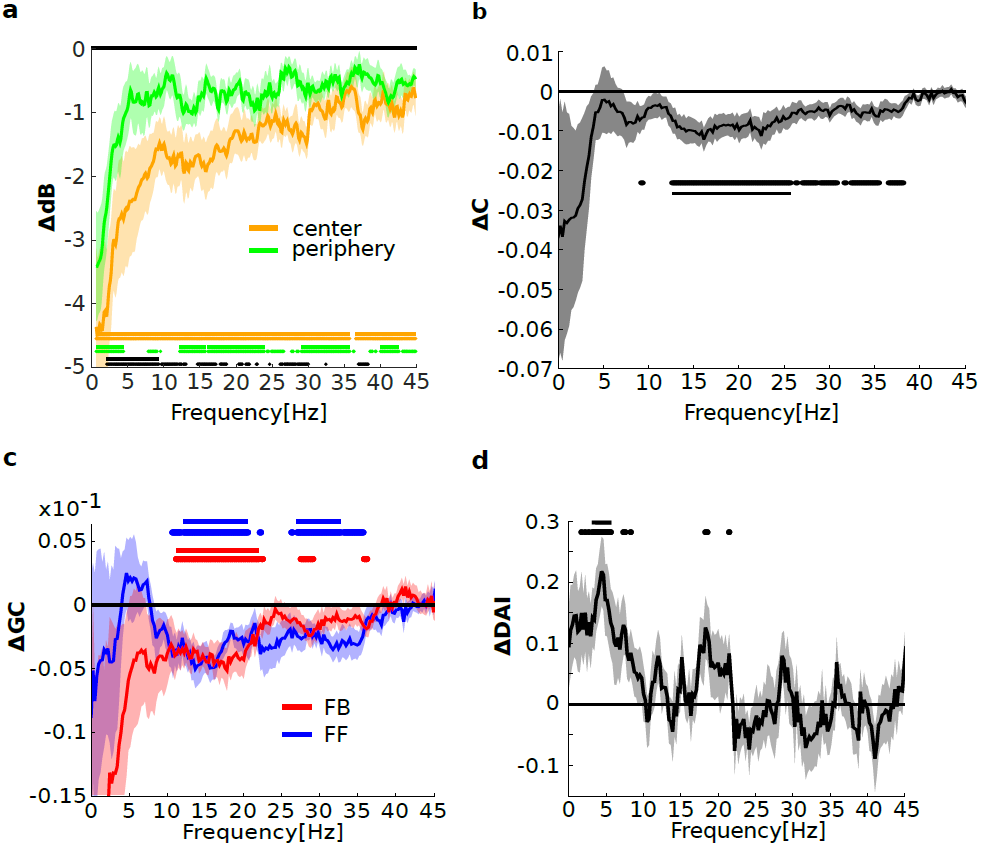
Isoflurane reduces power and coherence. **a)** The effect of isoflurane on center and periphery power, obtained by subtracting values in 0% isoflurane from values in 0.6% isoflurane. Negative values indicate that isoflurane reduced power for both the center and periphery. For central channels, orange dots and solid lines indicate uncorrected and corrected significant reductions due to anesthesia at the p< 0.05 level. For peripheral channels, green dots and solid lines indicate corresponding significant reductions at the p< 0.05 level. Black dots and solid line mark uncorrected and corrected, significantly greater reduction in the centre than the periphery at the p< 0.05 level. Shaded area represents sem across 13 flies. **b)** The effect of isoflurane on coherence, obtained by subtracting values in 0% isoflurane from values in 0.6% isoflurane. Black dots and solid line indicate uncorrected and correct significant reductions at the p< 0.05 level. Shaded area represent sem across 13 flies. **c)** The effect isoflurane on FF (blue) and FB (red) GC, obtained by subtracting values in 0% isoflurane from 0.6% isoflurane. Blue dots and solid line mark significant uncorrected and corrected reductions in FF GC at the p< 0.05 level. Red dots and line mark significant uncorrected and corrected reductions in FB GC at the p< 0.05 level. **d)** The effect of isoflurane on DAI of GC, obtained by subtracting values in 0% isoflurane from 0.6% isoflurane. Black dots and solid line mark significant uncorrected and corrected differences from zero at the p< 0.05 level

